# SpliceUp: Predicting *SF3B1* mutations in individual cells via aberrant splice site activation from scRNA-seq data

**DOI:** 10.1101/2025.09.22.677806

**Authors:** Maksim Kholmatov, Karin D. Prummel, Borhane Guezguez, Eva Hellström-Lindberg, Pedro L. Moura, Judith B. Zaugg

**Affiliations:** Molecular Systems Biology Unit, European Molecular Biology Laboratory (EMBL), Heidelberg, Germany; Department of Biomedicine, University Hospital Basel, University of Basel, Basel, Switzerland; Laboratory of Systems Biology and Genetics, Institute of Bioengineering, School of Life Sciences, École Polytechnique Fédérale de Lausanne (EPFL), Lausanne, Switzerland; Internal Medicine III, University Medical Center Mainz (UMC), Mainz, Germany; German Cancer Consortium (DKTK), Partner Site Frankfurt/Mainz, German Cancer Research Center (DKFZ), Heidelberg, Germany; Research Center for Immunotherapy (FZI), University Medical Centre of the Johannes Gutenberg University Mainz, Mainz, Germany; Center for Hematology and Regenerative Medicine, Department of Medicine Huddinge, Karolinska Institutet, Stockholm, Sweden

**Keywords:** *SF3B1*, myelodysplastic syndromes, splicing, single-cell

## Abstract

Myeloid neoplasms (MN) are clonal heterogeneous disorders initiated by somatic driver mutations in hematopoietic stem and progenitor cells (HSPCs). Among the most common are mutations in RNA splicing factors, which exert pleiotropic effects but are difficult to study due to altered hematopoietic differentiation and the inability to specifically isolate mutant HSPCs. Single-cell transcriptomics offers a powerful framework to dissect mutant cell states, yet direct genotyping is hampered by data sparsity and often requires costly, labor-intensive approaches. To overcome this limitation, we developed SpliceUp, a computational tool that identifies splice factor-mutant cells through their aberrant RNA splicing signatures. Applied to *SF3B1*-mutant samples, SpliceUp exploits cryptic 3’ splice site usage and exon-skipping events to achieve more than a 7-fold increase in mutant cell detection from low-coverage 10x Genomics datasets. Differential expression analysis revealed cell typespecific programs, including MHC-II upregulation in *SF3B1*-mutated HSCs and enhanced RNA translation in *SF3B1*-mutated erythroid precursors.

## Introduction

Myeloid neoplasms (MN) represent a heterogeneous group of clonal hematopoietic disorders, including myelodysplastic syndromes (MDS), myeloproliferative neoplasms (MPN), acute myeloid leukemia (AML), and related entities^1^. Despite their clinical diversity, these disorders share the common thread of selection and expansion of hematopoietic clones carrying somatic driver mutations^1^. Among the most recurrent drivers in MN are mutations affecting components of the spliceosome complex (splice factors), which produce characteristic and reproducible splicing defects in mature mRNA. For instance, *SF3B1* mutations promote activation of cryptic 3’ splice sites, whereas ZRSR2 mutations cause retention of U12-dependent introns^2^. Recent single-cell transcriptomic studies have further mapped missplicing in splice factor-mutated HSPCs, especially for *SF3B1* mutations, in MDS and clonal hematopoiesis, highlighting cell type-specific consequences of the mutation^3^. Their importance is underscored by their strong clonal fitness advantage even at low variant allele frequency (VAF)^4,5^, and their ability to drive specific disease-defining biology. Notably, *SF3B1* mutations frequently act as the primary, and sometimes sole, driver in MDS with ring sideroblasts (MDS-RS), a distinct subtype entity in the current WHO and IPSS-M classifications of MN^6,7^.

Assessing the specific transcriptomic consequences of MN driver mutations in bulk analysis is challenging, because altered differentiation skews cell composition and contaminating cells persist, even in CD34+ -HSPC-enriched material^8^. Single-cell approaches are therefore required. However, reliable identification of somatic mutations in scRNAseq remains challenging. With widely used platforms (10x Genomics, Drop-seq, Smartseq)^9–11^, direct identification of transcripts harboring the mutation is limited by data sparsity and low capture of mutated gene transcript regions. Moreover, biological variability driven by celltype identity often exceeds mutation-induced effects, complicating RNA expression-based detection strategies at single-cell resolution. Recent advances in experimental methods, including the implementation of targeted RNA/cDNA enrichment, can improve the sensitivity of mutation detection^3,12–14^; however, such methods remain labor-intensive, expensive, and/or dependent on access to the original sequencing libraries.

To address this gap, we developed SpliceUp (https://git.embl.org/grp-zaugg/spliceup), a novel computational framework that exploits mutation-associated splicing phenotypes to identify splice factor-mutant cells in scRNA-seq without additional experimental procedures. SpliceUp uses supervised learning on well-characterized aberrant splicing events, achieving a 7-fold increase in detection of mutant cells in sparse 10x Genomics datasets. By using *SF3B1*-mutant MDS samples as proof of principle, we demonstrate that SpliceUp is robust to data sparsity, scalable across datasets, and potentially generalizable toward the identification of cells with other recurrent splice factor mutations or known recurrent splicing errors.

## 2 Results

### Single-cell classification using alternative splicing events

SpliceUp classifies single cells into two classes (e.g., wild-type [WT] vs. splice factor-mutated [MUT]) based on a pre-defined set of differential splicing events (Figure 1A). The use of SpliceUp requires two inputs: (i) a set of paired splicing events distin-guishing the two cell classes and (ii) genome-aligned scRNA-seq data in the format of BAM files^16^ (Figure 1A). The cell-class specific splice-events, including skipped exons (SE), mutually ex-clusive exons (MXE), and alternative 3’/5’ splice sites (A3/5SS), can be obtained either from literature, public databases (e.g., MAJIQlopedia)^17^, or identified de novo from bulk or pseudo-bulk RNA-seq data using existing computational tools such as rMATS^18^, SUPPA2^19^, or MISO^20^.

**Figure 1.**
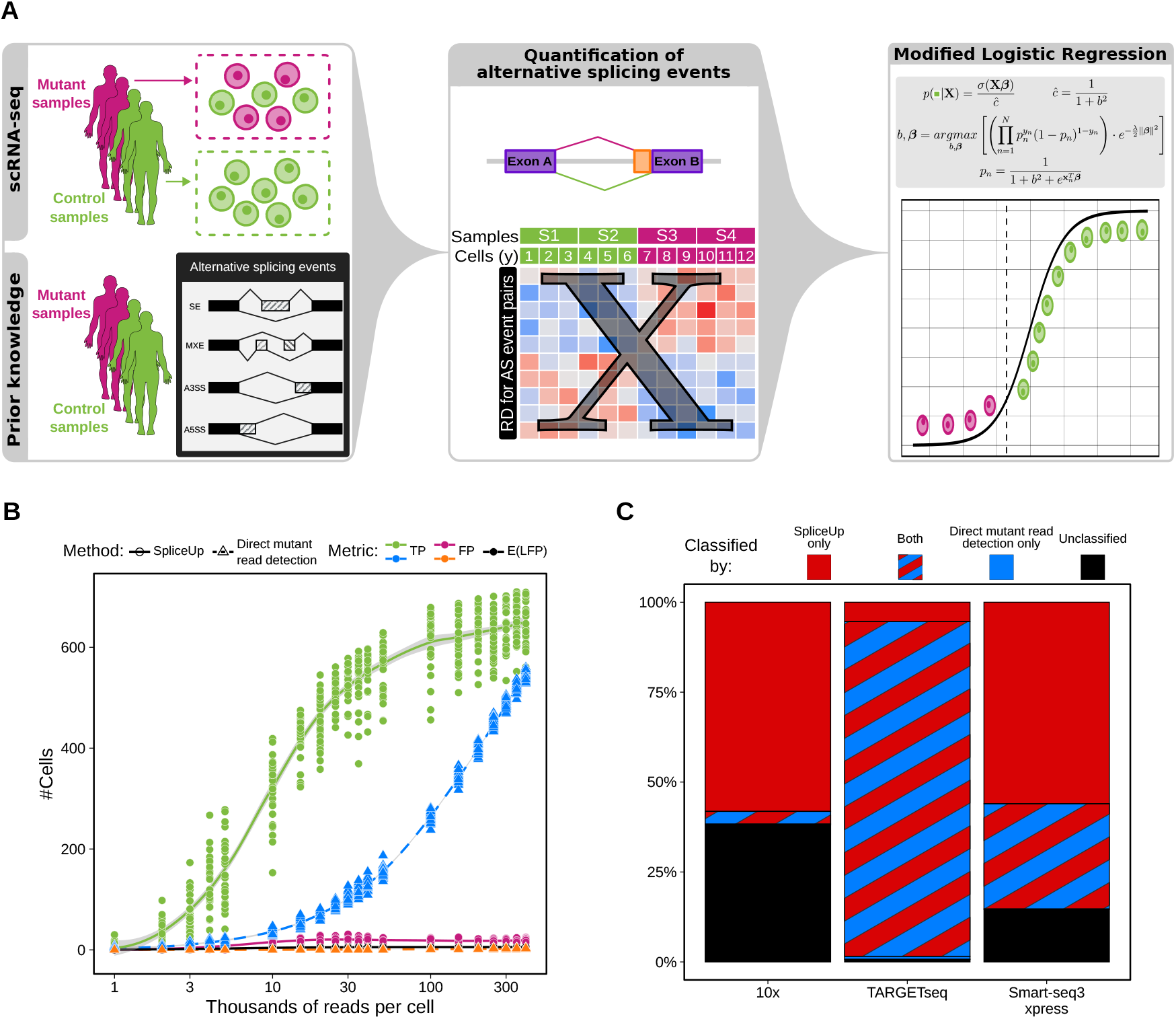
Description of the SpliceUp Method and benchmarking with TARGET-seq data. **A**) Conceptual description of SpliceUp: single-cell data comprises clean WT populations from control samples and mixed WT/MUT populations from patient samples. SpliceUp requires a table of miss-splicing events between WT and Mutants, quantifies the mis-splicing events in each cell, and fits a modified logistic regression model to calculate the probability of each cell being WT. **B**) Performance of SplicecUp against direct mutant read detection in TARGET-seq data from Moura et al. [15] at different sequencing depths. True positive (TP) and false positive (FP) predictions were defined based on PCR-based genotyping results. Additionally the expected number of labeled false positive (LFP) predictions coming from control samples was quantified based on the number of cells being classified and the FPR of 1% (Methods). **C**) Proportions of cells in each dataset used in this study that could be classified by either SpliceUp, direct mutant read detection or neither.

Based on these two inputs, SpliceUp quantifies the coverage for each of the pre-defined splicing event pairs (SEP) within each cell and calculates the relative difference (RD) between the two splice variants. As a final preprocessing step, splicing events that fail to distinguish the two classes on a pseudo-bulk level (user-defined threshold; default 5%) or have low coverage are removed (Methods). The resulting RD is combined in a cell-by-event-pair matrix that is then used as the independent variable for predicting the class (i.e., mutational status) of each cell.

A central challenge in training a single-cell classification model is the lack of reliable ground-truth labels, as patient samples typically contain both WT and mutant cells (Figure 1A). Partial labels for the MUT or WT groups can be obtained in two ways: cells from healthy controls can be assumed WT, and cells with sequencing reads covering the mutation site and confirming the mutant variant can be directly assumed MUT. However, in both cases, the opposite does not hold: mutant samples harbor residual WT cells, and the absence of mutant reads cannot be taken as confirmation of WT cell status, due to the limited sensitivity of single-cell sequencing. To overcome this challenge, we formulated the mutant cell classification as a Positive-Unlabeled (PU) learning problem: a binary classification task, where some observations from one class (by convention referred to as positive) are known, while the remaining observations are considered unlabeled and may belong to either class. When using sample labels, cells from the healthy control group serve as labeled positives (WT), while all other cells are considered as unlabeled. In contrast, when using direct mutation detection, cells with identified mutant reads are labeled as positives (MUT), and the rest remain unlabeled. A naive approach to this problem is considering all unlabeled cells as negatives and applying standard logistic regression (SLR). However, this approach is likely to overestimate the variance of the negative class, leading to misclassification of many true positives (Figure S1A).

To overcome these limitations of SLR we implemented a modified logistic regression (MLR) proposed by Jaskie, Elkan, and Spanias [21] (Methods). In addition to providing more accurate predictions in a PU-learning setting (Figure S1A), MLR explicitly estimates the probability that a positive observation is labeled as one of its parameters, *c*. This parameter can provide meaningful biological insight, and when an independent estimate exists (e.g., known VAFs based on existing clinical data), it offers an additional benchmark for model performance. Importantly, *c* should not be interpreted as the fraction of cells classified as positive; rather, it models the labeling process, whereas the final proportion of predicted mutant cells depends primarily on the choice of classification threshold (for example, a high *c* may coincide with relatively few classified mutant cells if a stringent threshold is applied).

### SpliceUp predicts *SF3B1* mutations of individual cells based on alternative splicing events

To benchmark SpliceUp’s performance, we assessed its ability to classify *SF3B1*-mutant cells from our previously published plate-based TARGET-seq^22^ dataset of HSPC subsets (HSC, hematopoietic stem cells, and MEP, megakaryocyte–erythroid progenitors) from three healthy controls and one MDS patient^15^. In this dataset, each cell’s ground truth label (WT or MUT) was established by PCR-based genotyping. We evaluated SpliceUp against a baseline approach that classified cells directly from mutant *SF3B1* transcript reads. The MDS sample contained two distinct *SF3B1*-mutant clones (K666N and N626D (Figure S1C, F)). To inform the choice of mutation-specific splicing events, we used a set of 648 events (319 SE, 192 A3SS, 49 A5SS, 88 MXE) previously identified by rMATS analysis of bulk RNA-seq from CD34^+^ HSPCs of *SF3B1*-K700E MDS patients and healthy donors^8^. SpliceUp was then trained using sample-level labels, designating WT as the positive class and MUT as unlabeled. Importantly, PCR-based genotyping was not used for model fitting, thereby allowing independent validation of predictions against ground-truth labels.

To perform predictions, we inverted the class labels along with predicted probabilities (Methods) and chose a classification threshold to limit the False Positive Rate (FPR) to 1% based on provided labels (Figure S1F, I). We note that the FPR estimate in the PU-learning setting is not equivalent to the real FPR calculated from ground-truth labels, but provides a practical approximation. Overall, SpliceUp achieved strong discriminatory power with AUC *≈* 0.96 (Figure S1K,I), with a real FPR of 3.2% (Table 1). Based on the estimated labeling probability, *c*, around 7.4% of the MDS cells were expected to be WT, which is a modest overestimation compared to the ground truth (5.2% WT).

We next assessed the performance of SpliceUp against the direct mutant read detection baseline across varying sequencing depths: for each method, we quantified the number of correctly classified MUT cells at different sequencing depths by randomly downsampling TARGET-seq reads per cell. Across the full range of coverages, SpliceUp consistently outperformed direct detection in recovering true positive predictions, albeit at the cost of a slightly higher FPR (Average FPR across all coverages *≈*3% for SpliceUp and *≈*0.2% for direct detection) (Figure 1B).

### SpliceUp identifies cell type-specific effects of *SF3B1* mutations

Next, we applied SpliceUp to dissect the cell type-specific effects of the *SF3B1* mutation in MDS patients. To this end, we analyzed BM cells from three healthy control samples, five *SF3B1*mutant MDS-RS samples, and additionally ring-sideroblasts enriched from two of the *SF3B1*-mutant MDS-RS donors profiled with 10x Genomics 3’ scRNA-seq^8^. In total, the data comprised 14,122 cells clustered into 12 cell types (Figure 2A, Figure S1D, Methods). Due to the inherent data sparsity of 3’ scRNA-seq, only *∼* 3.8% of cells contained reads overlapping the mutation site and could be classified by direct mutant read detection (Figure 1C), resulting in only *∼* 2.2% of cells directly classified as mutant (MUT). In contrast, SpliceUp predicted between 1.6% and 35.9% MUT cells across different BM cell types (Figure 2B, Figure S1G, J). In line with previous work^23^, very few MUT cells are predicted in plasma cells, stroma cells, and T cells, reflecting SpliceUp’s low false-positive rate. As expected from the MDS-RS phenotype, SpliceUp predicted that both HSPCs and erythroid progenitors harbored among the highest fractions of *SF3B1*-MUT cells (Figure 2B), consistent with their stem cell origin and the characteristic erythroid manifestation of the disease^24^. Notably, substantial fractions of B cells, monocytes, and NK cells (8-21%) were also predicted as MUT^23,25^.

**Figure 2.**
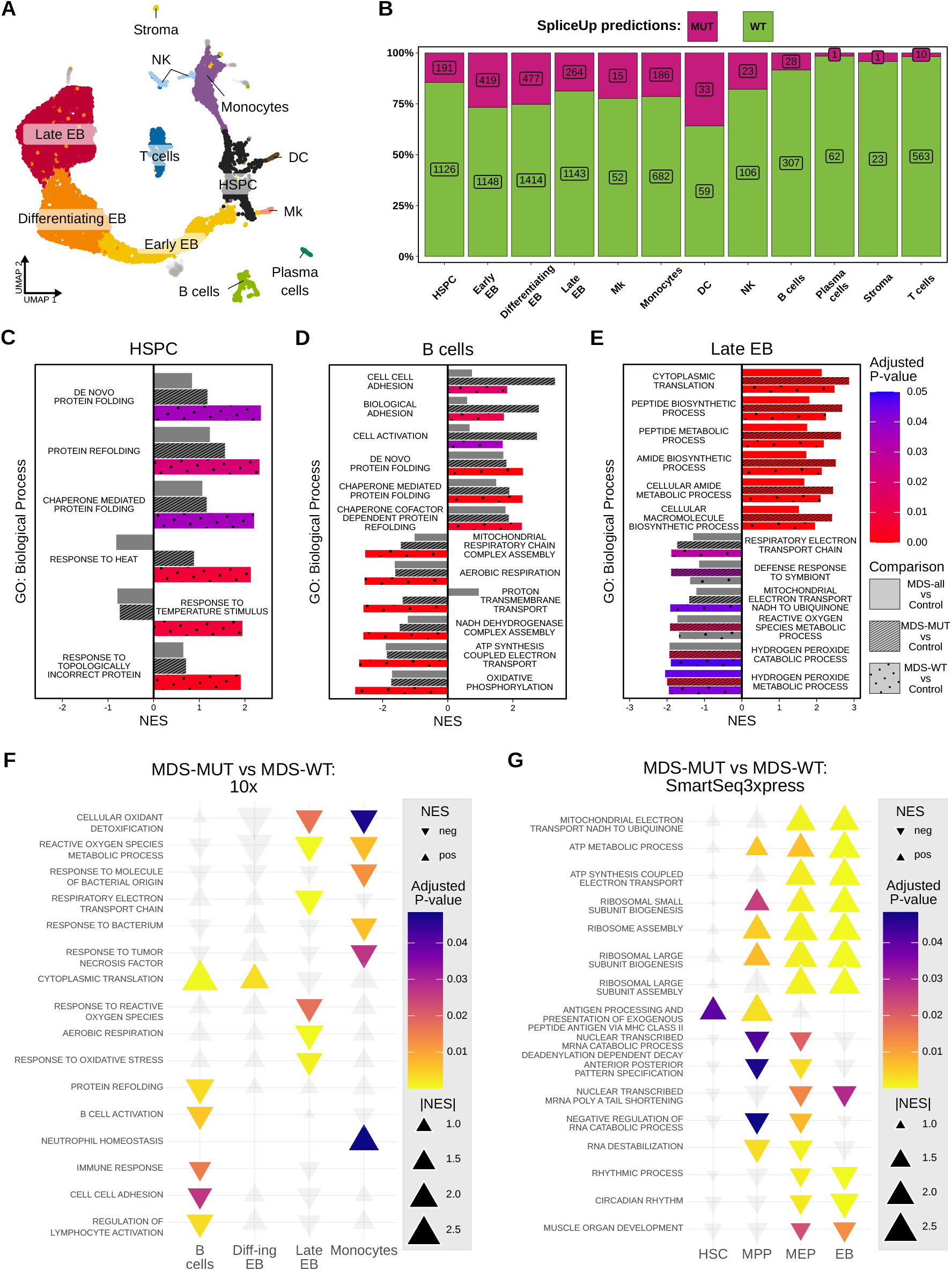
Identification of global disease-associated and mutation-intrinsic effects in MDS-*SF3B1*. Data from Moura et al. [8]: **A-F**) 10x dataset, **G**) SMART-seq3xpress dataset. **A**) UMAP representation of the 10x dataset colored based on the annotated cell types. **B**) Percentage of cells classified by SpliceUp as either MUT or WT within each cell type. Labels indicate the absolute number of cells in each group. Process collection. **C-E**) GSEA results with top positively and negatively enriched terms based on a pseudo-bulk comparison of MDS against control samples, using either all MDS cells (MDS-all), only MUT predictions from SpliceUp (MDS-MUT) or only cells classified as WT (MDS-WT) (Methods). Each of the cell types: **C**) HSPCs, **D**) B cells and **E**) Late EB; was analyzed independently. **F**) GSEA results with selected terms based on intra-patient comparison of SpliceUp-classified MUT and WT cells in the 10x dataset. For each cell type MUT and WT predictions across all MDS samples were compared in a pseudo-bulk fashion using sample identity as a covariate. **G**) GSEA results with top positively and negatively enriched terms based on single-cell comparison of SpliceUp-classified MUT and WT cells in the SMART-seq3xpress dataset. **C-G**) GSEA results against the terms from Gene Ontology: Biological Process collection. **C-E**) Positive values of NES (normalized enrichment score) signify enriched in MDS, negative NES in control. **F-G**) “pos” refers to enriched in the MUT over WT, “neg” refers to enriched in WT over MUT.

The ability of SpliceUp to classify cells into MDS-WT and MDS-MUT allowed us to compare each fraction separately to normal bone marrow, and thus also assess extrinsic effects of the MDS BM microenvironment on non-mutated (WT) cells. For each cell type, we quantified differential expression of the MDS-MUT and MDS-WT fractions separately against healthy controls (Figure S2), followed by gene set enrichment analysis (GSEA), which identified significantly enriched gene sets in HSPCs, B cells, and late erythroblasts (EBs) (Methods, Figure 2C-E). For HSPCs, we found the MDS-WT fraction enriched for heat-response and protein-folding pathways, largely driven by the upregulation of heat shock protein (HSP) genes (Figure 2C). These signatures were absent in the MUT-HSPCs in MDS, suggesting that *SF3B1*-mutant HSPCs engage less in stress-response programs or are intrinsically more buffered from proteotoxic stress, which could lead to their clonal advantage.

Similarly, the MDS-WT B cell fraction was enriched for protein folding and chaperone activity, alongside a downregulation of respiratory activity. Again, neither of these pathways was significantly enriched in the MUT fraction of MDS B cells (Figure 2D). This suggests that the altered pathways may reflect MDS BM microenvironmental effects on non-mutated B cells.

A different picture emerged for late EBs, where we found both MDS-MUT and MDS-WT fractions enriched for translation and RNA metabolism compared to healthy controls, indicating that increased translation is a general feature for late erythropoiesis in MDS (Figure 2E). At the same time, MDS EBs were depleted for aerobic respiration-related terms, consistent with previous observations on impaired aerobic respiration in erythroid cells in MDS-RS^8,26–28^ (Figure 2E). The observation that both fractions (MDS-MUT and MDS-WT) in EBs show similar enrichments compared to healthy EBs may reflect either a mutation-independent process driven by the diseased BM microenvironment or residual false-negatives among the predicted MDS-WT cells. Notably, when analyzed without separating MUT and WT fractions, MDS late EBs were the only cell type with significantly enriched pathways relative to healthy controls. However, these enrichments were weaker than those obtained using SpliceUp-classified fractions, indicating that SpliceUp improves detection of biologically relevant signals without the need to perform additional experimental protocols.

Next, we assessed the intrinsic effects of the *SF3B1*-mutation across the different cell types by comparing MDS-MUT against MDS-WT cells within each patient in a paired setting (Methods, Figure S3). We identified significantly enriched terms (GSEA) in B cells, differentiating EBs, late EBs, and monocytes. In B cells and differentiating EBs, we found upregulation of translation-related terms in the MDS-MUT, suggesting that upregulation of translation could be a general effect of the mutation (Figure 2F, Figure S4). In contrast, MDS-WT B cells were enriched for terms related to B cell activation, indicating that B cells that harbor the mutation are less functional. For the monocytes, we found MDS-WT enriched for processes associated with responses to cytokines, vascular growth factor production, response to bacterial signals, TNF, and oxidative stress response, while MDS-MUT were enriched for neutrophil homeostasis, which could hint at a role of the *SF3B1*-mutation in altering monocyte function.

Finally, we further investigated the different HSPC populations using an additional data set of FACS-purified CD34^+^ HSPCs of one *SF3B1* (K700E)-MDS patient, deeply profiled with SmartSeq3-xpress^29^ from Moura et al. [8] (Figure S1E). The variant allele frequency (VAF) was around 32%, corresponding to *∼* 64% of cells expected to be mutant. The baseline (direct mutant read detection) identified *∼* 15% (251/1650) *SF3B1*-K700E mutant cells based on direct coverage of the mutated transcript (Figure S1H), which, due to SMART-seq’s full-length transcript capture and the higher sequencing depth was substantially higher than in the 10x dataset. This allowed us to apply SpliceUp by using the mutant cells recovered with direct mutant read detection as a positive-labeled class (Methods, Figure S1K). SpliceUp predictions provided an additional 424 (total 675) mutant cells (*∼* 48%), which is closer to what we expect from the VAF (Figure 1C).

Additionally, mutated MPPs showed enrichment for translational terms (e.g., ribosome assembly), which were even more pronounced in MEPs and EBs. These findings align with our observations in the 10x dataset. While the translational hyperactivity of *SF3B1*-mutated MDS cells has been described before^30^, our results reveal an additional and not yet described feature: Both *SF3B1*-mutant HSC and MPP cells display increased antigen presentation and MHC-II activity (Figure 2G). This novel insight is in line with recent findings in AML, where leukemic stem cells exhibit increased MHC-II activity^31^ and suggest that aberrant antigen presentation may represent a shared vulnerability of mutant HSPCs across myeloid malignancies.

Together, these analyses demonstrate the power of SpliceUp to resolve and identify relevant biological processes in the MDS BM microenvironment, both in a small patient cohort, and in mutated cells from a single patient sequenced at high depth.

## 3 Discussion

This study presents SpliceUp, a novel computational approach for identifying splice factor-mutated cells in single-cell transcriptomic datasets. We validate SpliceUp against PCR-based genotyping in a dataset containing *SF3B1*-mutant cells. We demonstrate that SpliceUp yields biologically meaningful predictions that can improve the performance of downstream analyses by increasing the statistical power of differential expression studies in samples composed of mixtures between WT and mutant cells. Compared to direct mutant-read-based classification, SpliceUp improves the sensitivity of mutation detection. While it is unlikely to outperform experimental methods specifically designed for single-cell genotyping (e.g., GoT or RaCH-seq), it uniquely enables in-depth investigation of mutation-associated transcriptional changes from scRNA-seq data without the need of additional experimental assays or access to the original sequencing libraries. This makes SpliceUp particularly valuable in the current era where droplet-based scRNA-seq has become routine, as it runs on data produced with the most common single-cell technologies without the complexity of producing and integrating multiple specialized datasets.

In principle, SpliceUp can be applied to detect a broad range of splice factor mutations that give rise to distinct mis-splicing patterns, including recurrent lesions in *SRSF2* and *U2AF1* that are quite common in MDS but also in other MN and blood malignancies. By improving the computational identification of somatic mutations directly from high-throughput transcriptomic data, SpliceUp facilitates more powerful comparisons of malignant versus healthy counterparts and enables deeper investigations into tumor biology, including intratumoral heterogeneity, clonal architecture, and growth dynamics.

In conclusion, SpliceUp provides a broadly applicable framework for linking splicing factor mutations to their cell-type–specific transcriptional consequences, thereby extending the utility of existing and future single-cell transcriptomic datasets in hematologic malignancies and beyond.

### Limitations of the study

Several factors may affect the performance of SpliceUp. First, the method relies on the availability of a high-quality reference of alternative splicing events specific to the mutational or disease context being predicted, which can be limiting if no such reference is publicly accessible. Second, the modified logistic regression (MLR) classification model is based on several assumptions that may not hold under all biological contexts. In par-ticular, the SCAR assumption (selected completely at random) in the setting of mutation prediction implies that WT cells from diseased conditions exhibit splicing patterns comparable to those in healthy tissues. In reality, cells in the tumor microenvironment are frequently functionally altered by interactions with malignant and immune cells, and these effects are reflected in their RNA expression^32^. Some effects of hypoxia and pro-inflammatory cytokine signalling on alternative splicing in non-cancer tissues have been described in the context of ovarian cancer^33,34^ and pancreatic islets^35^, but the extent to which such interactions specifically affect alternative splicing in the bone marrow context and their overlap with mutation-associated mis-splicing events remains poorly understood. Violation of the SCAR assumption would lead to reduced model performance and limit the ability to control the false positive rate (FPR) when setting a classification threshold. Recently some PU-learning approaches that do not rely on the SCAR assumption or relax it have been proposed^36–38^. Adapting and applying these methods to mutation prediction in single-cell data represents a promising avenue for future research.

## 4 Materials and Methods

## Single-cell data source

TARGET-seq data from Moura et al. [15] was obtained from the Swedish National Data Service’s research data repository: SND-ID: 2023-223. 10x single-cell RNA-seq and Smartseq3xpress data from Moura et al. [8] was obtained from the Swedish National Data Service’s research data repository: SNDID 2023–121, SND-ID 2023–122.

All source material was provided with written informed consent for research use, given in accordance with the Declaration of Helsinki, and the study was approved by the Ethics Research Committee at Karolinska Institutet (2022-03406-02, 2024-03119-02).

### Quality control of single-cell RNA-seq data

Downstream analysis of single-cell data were performed using *Seurat* v4.3.0.1^39^ unless otherwise specified.

#### TARGET-seq

Count matrix and metadata were converted to a SeuratObject^39^. Percent of transcripts mapping to mitochondrial genes were calculated using *Seurat*’s *PercentageFeatureSet* function. Cells with < 800 genes detected and > 15% transcripts mapping to mitochondrial genes were removed.

#### Smart-seq3xpress

Count matrix based on exon and intron mapped reads and metadata was converted to a SeuratObject. Cells with < 1000 genes detected and < 1000 UMIs were removed.

#### 10x

Pre-processed Seurat object was obtained from the authors of Moura et al. [8].

### Normalization and dimensionality reduction

TARGET-seq and Smart-seq3xpress data were log-normalized (*NormalizeData*) and 2000 most highly variable genes were selected (*FindVariableFeatures*). Matrix of variable gene expression was scaled and centered (*ScaleData*) and used as input for PCA (*RunPCA*). First 30 principal components were used to perform UMAP dimensionality reduction (*RunUMAP*). The K-nearest neighbor (kNN) graph was constructed based on the first 10 principal components (*FindNeighbors*).

### Sub-clustering and cell type annotation

The 10× dataset from Moura et al. [8] contained a cluster of cells not characterized in the original publication, that consisted of several non-myeloid population and potential clusters of doublets (Figure S1D). We additinally sub-clustered this group of cells using the *FindSubCluster* function from *Seurat* with the Leiden algorithm^40^ *resolution = 0*.*1, algorithm = 4*. Among the individual sub-clusters we identified Stromal, B cell, Plasma cell and NK cell populations (Figure 2A) based on known marker genes identified using *Seurat*’s *FindAllMarkers* function.

### Quantification of splicing event support

Splicing event quantification was performed using a custom fork^41^ of pysam^42,43^. To quantify the support for each splicing event we select all reads overlapping the corresponding splice site coordinates from the BAM file and check for the presence of the N operation in CIGAR string immediately adjacent to the splice site, which for mRNA-to-genome alignment represents the presence of an intron as defined in the SAM format specification^16^. For all such reads we record the corrected cell barcode sequences and corrected UMI sequences, if available, and quantify the number of UMIs/reads per splice site per cell barcode.

### Filtering of alternative splicing events

To exclude alternative splicing event pairs that are not covered in single-cell data we aggregate their average coverage per sample and exclude events where both alternatives have average coverage lower than 1% in 75% of samples or more. With reasonably high numbers of cells from the mutant clone we expect the mutation-specific events to be able to separate our samples in a pseudo-bulk comparison (Figure S1B). To exclude alternative splicing event pairs that don’t differ between conditions we calculate relative difference (*RD*) between average coverage for alternative 1 (*A*_1_) and alternative 2 (*A*_2_) within each group:

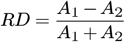

and a fractional relative difference (*FRD*) between the groups (*g*1, *g*2) for each splicing event pair:

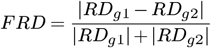

We exclude the splicing event pairs that show FRD below a threshold of 5%.

### Imputation of missing values

Due to the sparsity of single-cell data some cells will have no reads supporting either of the alternatives for a given splicing event pair, resulting in missing values. To be able to fit a classification model we impute these missing values by either using the average across the non-missing values or using predictive mean matching via the *mice* R package^44^. In our experience using the average to for imputation worked well in general, except for datasets with highly imbalanced class distributions like Smart-seq3xpress where predictive mean matching has significantly improved model’s performance.

### Modified and standard logistic regression

To perform the classification of cells based on their mutational status we implemented standard logistic regression approach and modified logistic regression method described in Jaskie, Elkan, and Spanias [21] as part of the *PuMLR* R package. In SLR we model the probability of an observation *i* belonging to the positive class as:

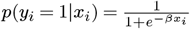

Parameter estimation is done by minimizing the total cross-entropy loss across observations.

MLR method makes two assumptions about the data. First is the partial separability assumption: the region of highest density of the labeled-positive observations consists entirely of positive observations. Second is the selected completely at random (SCAR) assumption: labeled-positive observations are selected completely randomly from the set of all positive observations. Based on these assumptions MLR consists of a two-step procedure. First step consists of learning of a non-traditional classifier for the observation being labeled. Probability of an observation being positive-labeled is modeled as:

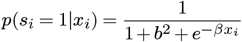

where parameter *b* creates an upper bound on the predicted probabilities. The value of the upper bound is related to *b* as follows:

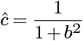

The second step utilizes the estimated *c* to estimate the probability of an observation coming from a positive class as:

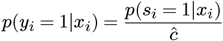

Parameters are estimated by minimizing the total cross-entropy loss across observations with an additional small ridge regularization (*λ* = 0.01) for improved conditioning and convergence (Figure 1A).

Loss functions for SLR and MLR are minimized using the Low-storage BFGS algorithm^45^ implemented in the *NLopt* library^46^.

### Mutation prediction in single-cell data

#### TARGET-seq, 10x

For training the MLR model we assign the sample level mutational status to all cells. WT cells (i.e. cells from control samples) were encoded to be positive-labeled class (*s* = 1) and MDS cells were encoded to be unlabeled class (*s* = 0). SEP’s were filtered as described above and cells with < 4 SEP covered by sequencing reads were excluded from the training. Missing values were imputed using the avarage RD across all cells. Resulting cell-bySEP matrix was used as a predictor (*X*) to fit the MLR model as described above. Predicted positive class probabilities (*p*) were inverted (1*− p*) and used to define the classification threshold by measuring FPR at different threshold values using the performance function from the *ROCR* (v1.0.11)^47^ R package. Final classification threshold was chosen as the largest cut-off for which FPR *<*= 1%. The sample enriched for ring-sideroblasts in 10x data showed substantially lower recovery of mutant cells despite the enrichment, which could be related to overall lower data quality, and therefore was excluded from downstream differential expression and GSEA.

#### Smart-seq3xpress

Mutational status derived from direct mutant read detection was applied as a training label for the MLR model with mutant cells encoded to be positive-labeled class (*s* = 1) and all other cells as unlabeled class (*s* = 0). Cells with < 4 SEP covered by sequencing reads were excluded from the training. Missing values in the RD matrix were imputed using predictive mean matching. Classification threshold was chosen to optimize the F1 score, calculated using the *ROCR* (v1.0.11)^47^ R package.

### Differential gene expression and GSEA analyses

#### Single-cell analysis of Smart-seq3xpress data

Log2 fold changes were estimated in each cell type using the *MAST* (v1.24.1)^48^ method via *Seurat*’s *FindMarkers* function.

#### Pseudo-bulk analysis of 10x data

Raw counts from individual cells were aggregated based on either the sample status or a combination of a sample identity (for control group) and predicted mutation status (for MDS samples) using *Seurat*’s *AggregateExpression* function. Only genes with *≥* 10 aggregated counts in at least 3 samples were kept. Aggregated counts were analyzed for differential expression in one of the settings describeg bellow using the *DESeq2* (v1.38.3)^49^ package. P-values were adjusted using the *IHW* ^50^ method. Log2 fold change values resulting from each analysis were shrunken using *apeglm* (v1.20.0)^51^ method (*lfcShrink(type=“apeglm”)*). The analyses in Figure 2C-E were done between different subsets of MDS cells and control sample cells. Comparison of MDS-MUT cells vs controls was done by only keeping cells classified as mutant by SpliceUp in MDS samples. Selection of MDS-WT cells is much more challenging, because while we can approximate the FPR by using cells from control samples as reliably labeled WT, we have no way of measuring the FNR. In lieu of any way to control for the contamination of the WT population with MUT cells, we applied a more stringent classification threshold within each cell population based on probabilities predicted with SpliceUp and chose a value that maximizes the downstream biological signal (the number of significantly enriched biological processes). The final thresholds for selecting the WT population were: 15% for HSPCs and 50% for B cells and Late EBs. The comparison of MUT and WT predictions in Figure 2F was performed within MDS samples using aggregated counts per cell type per sample, split into MUT and WT groups by applying the standard SpliceUp threshold based on FPR control, with the sample identity used as a covariate to account for between-sample variability.

#### GSEA

Gene ontology Biological Process gene sets^52,53^ were obtained using the *msigdbr* (v7.5.1)^54^ R package. Gene set enrichment analysis was performed using the *GSEA* function from the *clusterProfiler* (v4.6.2)^55^ package with “nPerm=1e6” and “pvalueCutoff=1”.

## 5 Code availability

Code for the SpliceUp method is available at: https://git.embl.org/grp-zaugg/spliceup. Implementation of the Modified Logistic Regression and traditional Logistic regression is available as an R package -PuMLR available at https://github.com/maxim-h/PuMLR. Code reproducing the analysis performed will be made available upon publication of this manuscript.

## AUTHOR CONTRIBUTIONS

MK, PLM and JBZ conceived and designed the study. MK developed the SpliceUp method and performed the data analysis under supervision of PLM, JBZ and EHL. MK, KDP, BG, PLM and JBZ wrote the manuscript with input from EHL.

## ACKNOWLEDGEMENTS

MK is funded by the EC H2020 MSCA-ITN Project ENHPATHY (grant agreement number 860002) to JBZ. KDP is supported by the SNSF (P2ZHP3_199669) and EMBO (538-2021) Postdoctoral Fellowships. BG is supported by the German Cancer Consortium (DKTK) Joint Funding Program (DKTK CHOICE). PLM is supported by the Myelodysplastic Syndromes Foundation, Inc. (grant number 1142079), the European Hematology Association (Topic-In-Focus Advanced Research Grant 2025), the Dr. Åke Olsson foundation (Dnr 2024-00303), the Alex and Eva Wall-ström foundation (Dnr 2024-00311) and a KI Research Foundation grant (Dnr 2024-02330). JBZ is supported by the European Union (ERC, epiNicheAML, 101044873).

Views and opinions expressed are however those of the authors only and do not necessarily reflect those of the European Union or the European Research Council. Neither the European Union nor the granting authority can be held responsible for them.

