## Supplementary Figures for "SpliceUp: Predicting *SF3B1* mutations in individual cells via aberrant splice site activation from scRNA-seq data"

### Supplementary Information

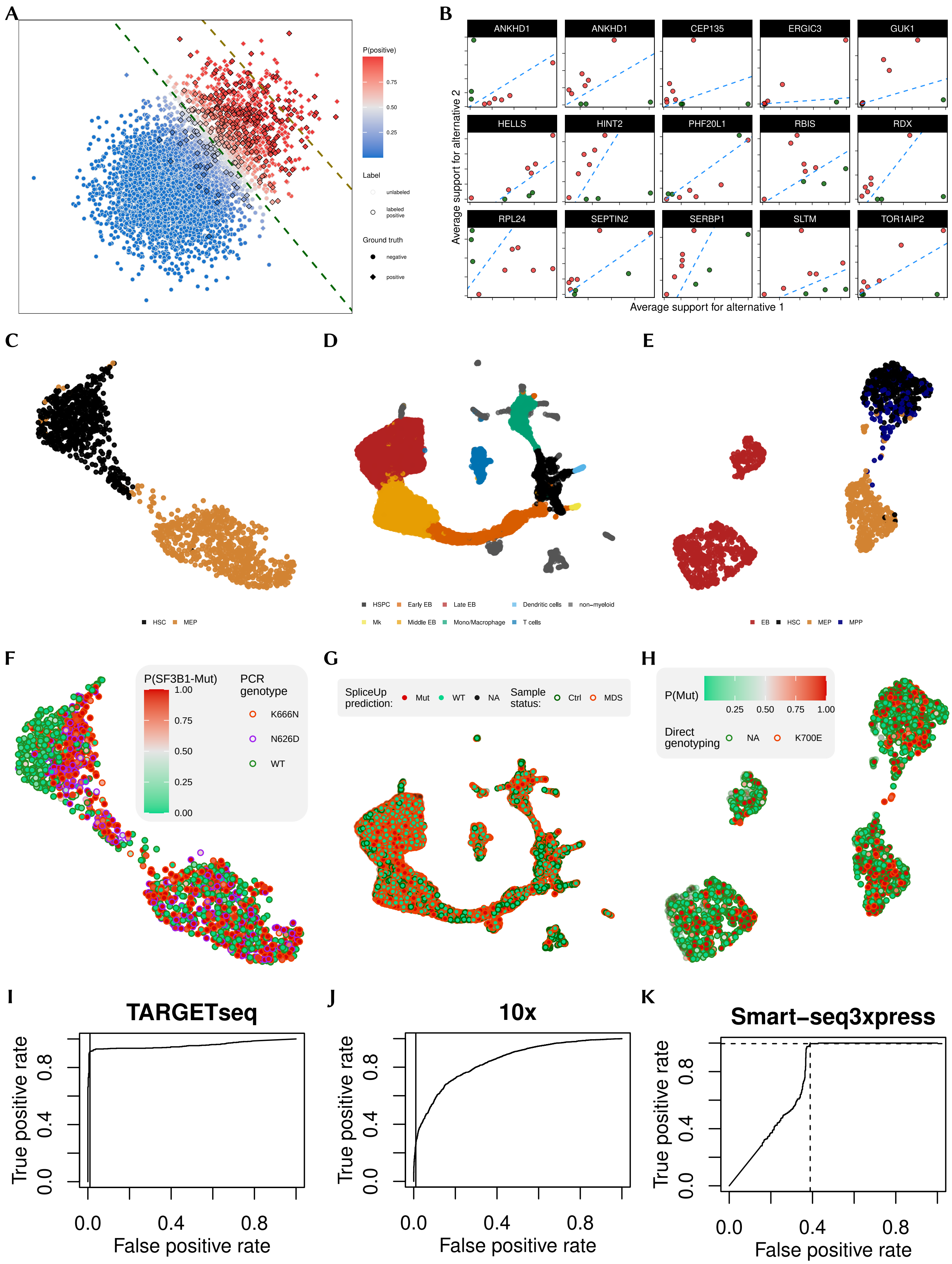
A
P(positive)
0.75
0.50
0.25
Label
unlabeled
labeled positive
Ground truth
negative
positive
B
ANKHD1
ANKHD1
CEP135
ERGIC3
GUK1
HELLS
HINT2
PHF20L1
RBIS
RDX
RPL24
SEPTIN2
SERBP1
SLTM
TOR1AIP2
Average support for alternative 2
Average support for alternative 1
C
D
E
HSC
MEP
HSPC
Early EB
Late EB
Dendritic cells
non−myeloid
Mk
Middle EB
Mono/Macrophage
T cells
EB
HSC
MEP
MPP
F
G
H
P(SF3B1-Mut)
1.00
0.75
0.50
0.25
0.00
PCR genotype
K666N
N626D
WT
SpliceUp prediction:
Mut
WT
NA
Sample status:
Ctrl
MDS
P(Mut)
0.25
0.50
0.75
1.00
Direct genotyping
NA
K700E
I
J
K
TARGETseq
10x
Smart−seq3xpress
True positive rate
False positive rate
0.0
0.4
0.8

**Figure S1. Overview of MLR, alternative splicing in MDS, dataset composition and predictive performance of SpliceUp**
**A**) PU-learning problem example with simulated data. Point color indicates positive class probabilities predicted with MLR. Dashed lines indicate the decision boundaries at 50% probability for MLR (green) and SLR (yellow). **B**) Pseudo-bulk coverage of alternative splicing events that can fully discriminate control and MDS-RS samples from Moura et al. [8]. Dashed lines indicate the decision boundary. **C-E**) UMAP of 3 datasets used in this study: TARGET-seq (**C**), 10x (**C**) and Smart-seq3xpress (**D**) colored by the cell type annotated by original authors. **F-G**) UMAP of 3 datasets used in this study colored by the labels used to fit the classification model by SpliceUp: for TARGET-seq and 10x (**F**, **G**) labels are based on the sample status. For Smart-seq3xpress (**H**) labels are based in the direct detection of mutant reads associated with each cell barcode. **I-K**) ROC curves based on the SpliceUp predictions within each dataset. Vertical lines in (**I**, **J**) represent the pre-defined FPR cut-off of 1% in TARGET-seq and 10x data. Vertical and horizontal lines in (**K**) represent the values corresponding to the maximum F1 score, which was used do define the classification threshold.

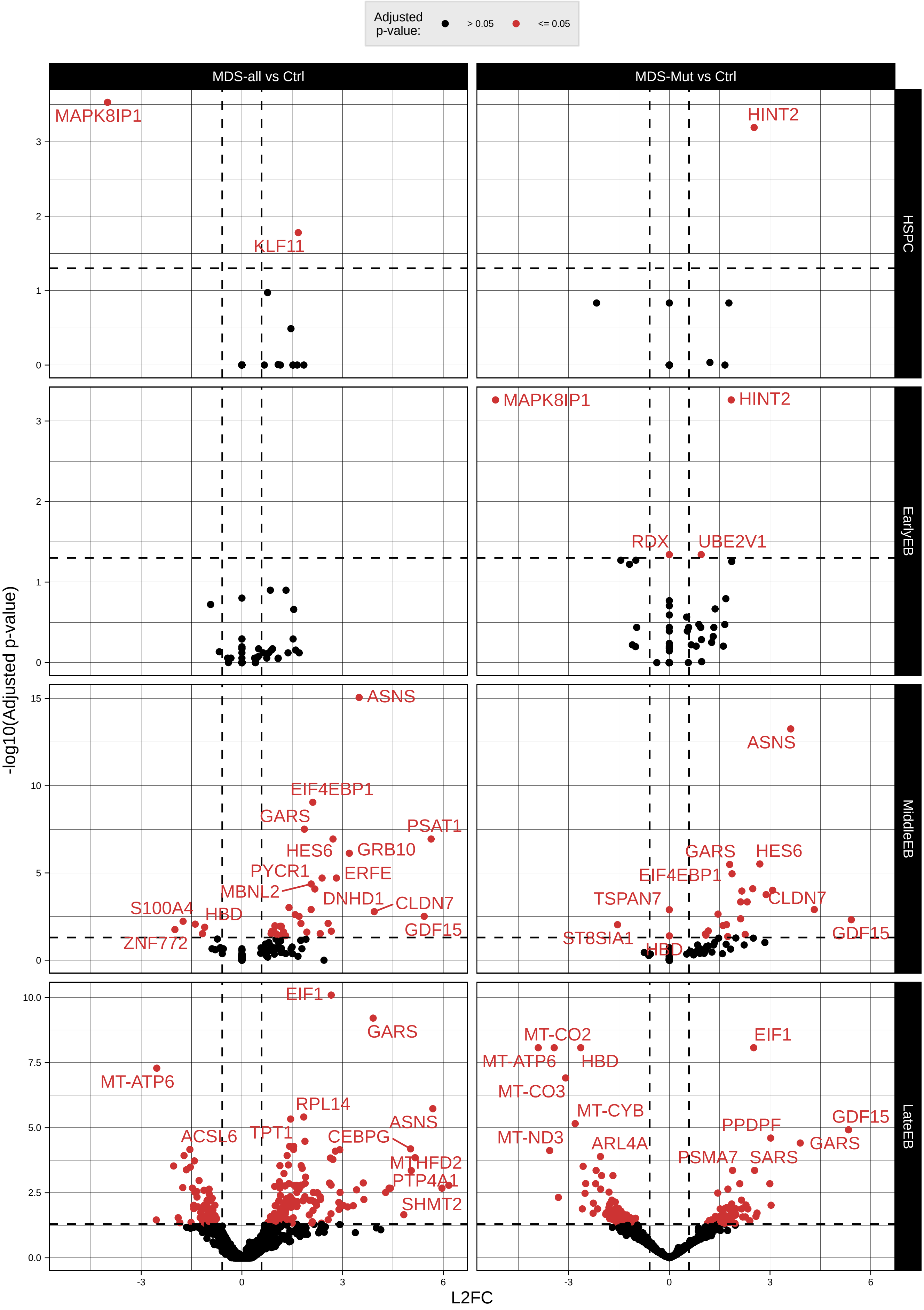
Adjusted p-value:
> 0.05
<= 0.05
MDS-all vs Ctrl
MDS-Mut vs Ctrl
HSPC
EarlyEB
MiddleEB
LateEB
-log10(Adjusted p-value)
L2FC
MAPK8IP1
KLF11
HINT2
RDX
UBE2V1
ASNS
EIF4EBP1
GARS
PSAT1
HES6
GRB10
PYCR1
ERFE
MBNL2
DNHD1
S100A4
HBD
CLDN7
GDF15
ZNF772
TSPAN7
ST8SIA1
EIF1
MT-ATP6
RPL14
ACSL6
TPT1
CEBPG
MTHFD2
PTP4A1
SHMT2
MT-CO2
MT-CO3
MT-CYB
MT-ND3
ARL4A
PPDPF
PSMA7
SARS

**Figure S2. Differential expression between MDS and control samples**
Volcano plots showing differential expression results for cell types with at least 1 significant gene. Left column represents the baseline pseudo-bulk comparison of all cell in a given cell type between MDS and control samples. The right column shows the same analysis, but only including MDS cells classified as MUT.

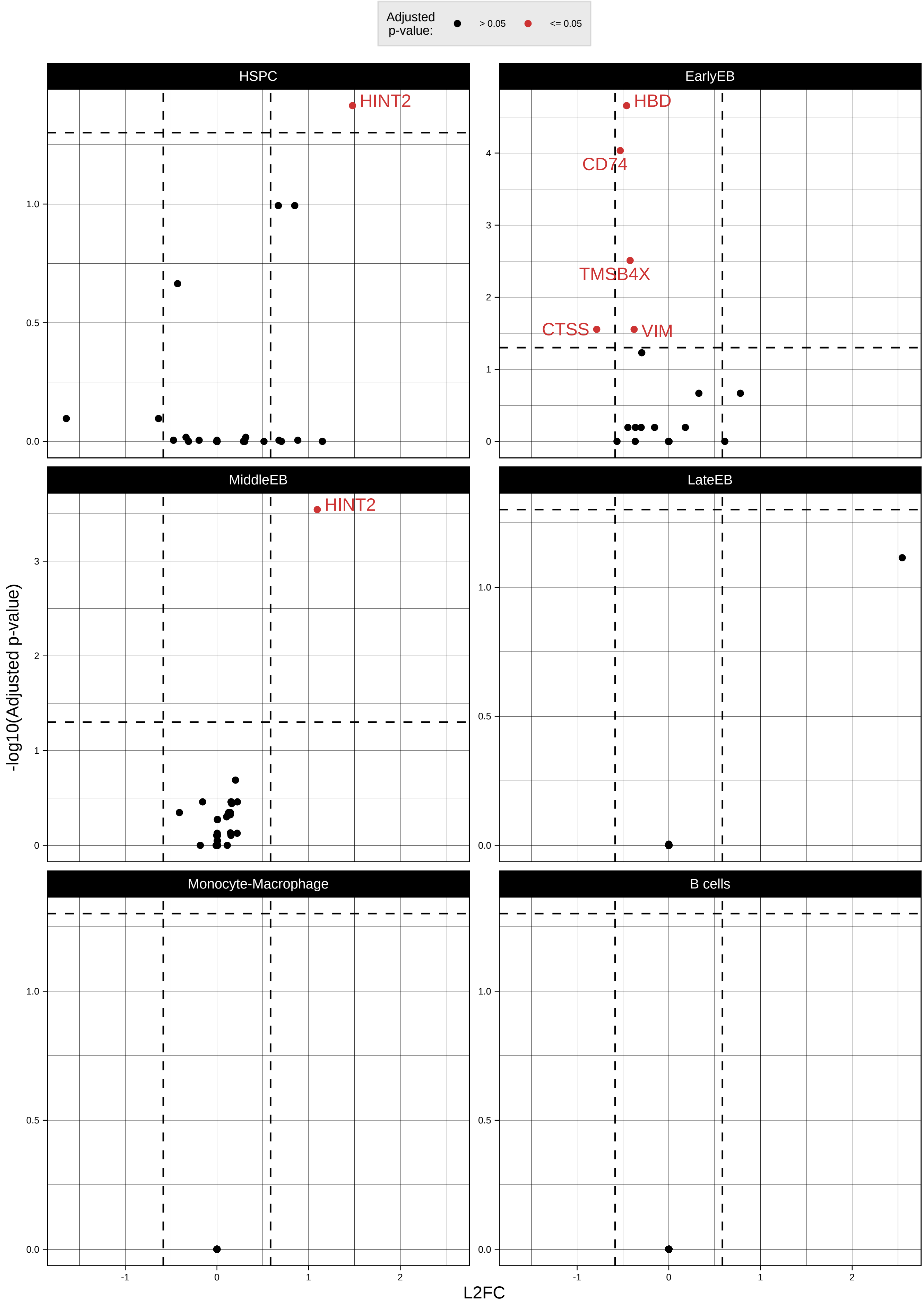
Adjusted p-value:
> 0.05
<= 0.05
HSPC
EarlyEB
MiddleEB
LateEB
Monocyte-Macrophage
B cells
HINT2
HBD
CD74
TMSB4X
CTSS
VIM
HINT2
-log10(Adjusted p-value)
L2FC

**Figure S3. Mutation-intrinsic differential expression in MDS samples**
Volcano plots showing differential expression results for the pseudo-bulk paired comparison of cells classified as MUT and WT in MDS samples by SpliceUp.

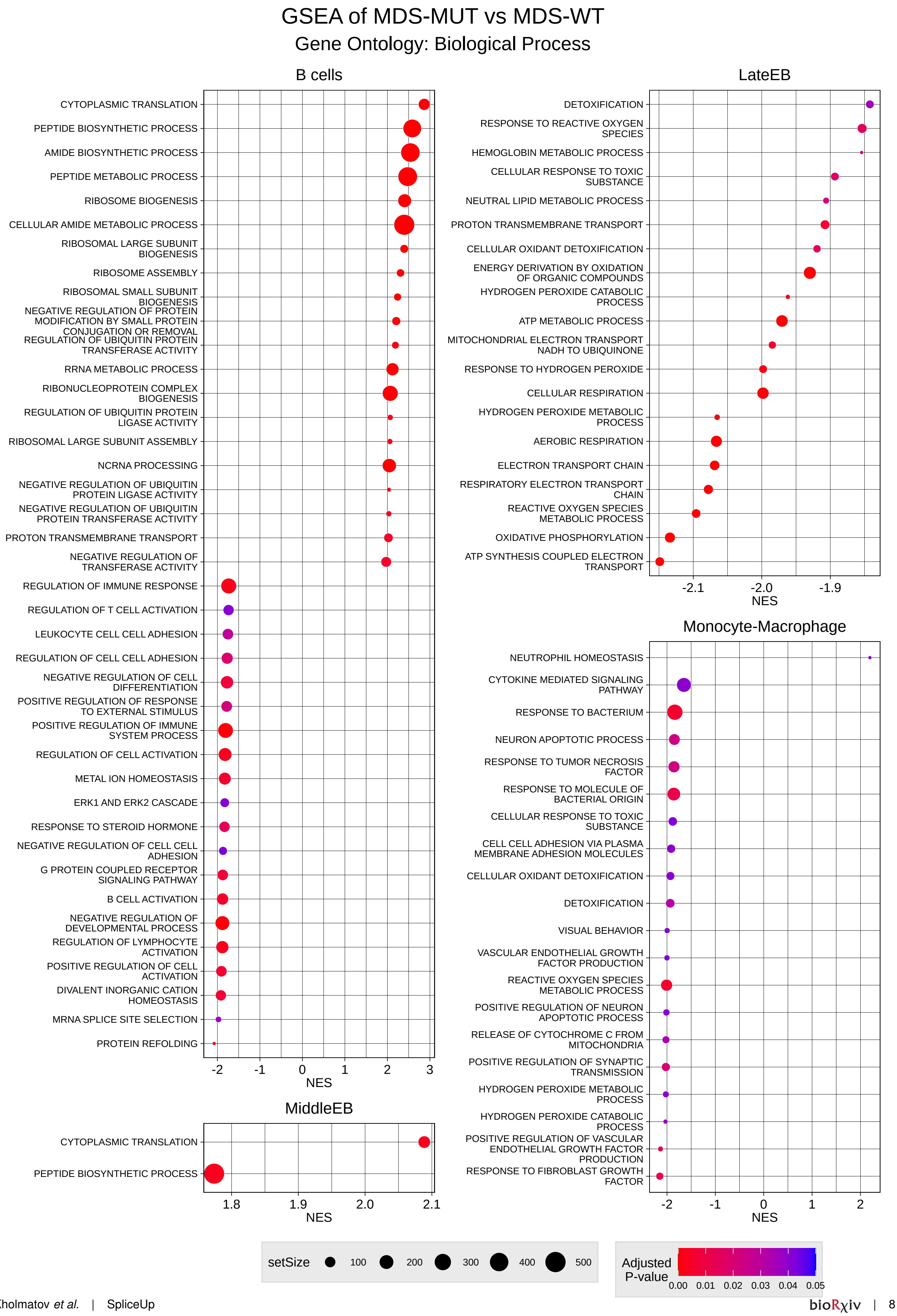

GSEA of MDS-MUT vs MDS-WT
Gene Ontology: Biological Process
B cells
CYTOPLASMIC TRANSLATION
PEPTIDE BIOSYNTHETIC PROCESS
AMIDE BIOSYNTHETIC PROCESS
PEPTIDE METABOLIC PROCESS
RIBOSOME BIOGENESIS
CELLULAR AMIDE METABOLIC PROCESS
RIBOSOMAL LARGE SUBUNIT BIOGENESIS
RIBOSOME ASSEMBLY
RIBOSOMAL SMALL SUBUNIT BIOGENESIS
NEGATIVE REGULATION OF PROTEIN MODIFICATION BY SMALL PROTEIN CONJUGATION OR REMOVAL
REGULATION OF UBIQUITIN PROTEIN TRANSFERASE ACTIVITY
RRNA METABOLIC PROCESS
RIBONUCLEOPROTEIN COMPLEX BIOGENESIS
REGULATION OF UBIQUITIN PROTEIN LIGASE ACTIVITY
RIBOSOMAL LARGE SUBUNIT ASSEMBLY
NCRNA PROCESSING
NEGATIVE REGULATION OF UBIQUITIN PROTEIN LIGASE ACTIVITY
NEGATIVE REGULATION OF UBIQUITIN PROTEIN TRANSFERASE ACTIVITY
PROTON TRANSMEMBRANE TRANSPORT
NEGATIVE REGULATION OF TRANSFERASE ACTIVITY
REGULATION OF IMMUNE RESPONSE
REGULATION OF T CELL ACTIVATION
LEUKOCYTE CELL CELL ADHESION
REGULATION OF CELL CELL ADHESION
NEGATIVE REGULATION OF CELL DIFFERENTIATION
POSITIVE REGULATION OF RESPONSE TO EXTERNAL STIMULUS
POSITIVE REGULATION OF IMMUNE SYSTEM PROCESS
REGULATION OF CELL ACTIVATION
METAL ION HOMEOSTASIS
ERK1 AND ERK2 CASCADE
RESPONSE TO STEROID HORMONE
NEGATIVE REGULATION OF CELL CELL ADHESION
G PROTEIN COUPLED RECEPTOR SIGNALING PATHWAY
B CELL ACTIVATION
NEGATIVE REGULATION OF DEVELOPMENTAL PROCESS
REGULATION OF LYMPHOCYTE ACTIVATION
POSITIVE REGULATION OF CELL ACTIVATION
DIVALENT INORGANIC CATION HOMEOSTASIS
MRNA SPLICE SITE SELECTION
PROTEIN REFOLDING
-2 -1 0 1 2 3
NES
LateEB
DETOXIFICATION
RESPONSE TO REACTIVE OXYGEN SPECIES
HEMOGLOBIN METABOLIC PROCESS
CELLULAR RESPONSE TO TOXIC SUBSTANCE
NEUTRAL LIPID METABOLIC PROCESS
PROTON TRANSMEMBRANE TRANSPORT
CELLULAR OXIDANT DETOXIFICATION
ENERGY DERIVATION BY OXIDATION OF ORGANIC COMPOUNDS
HYDROGEN PEROXIDE CATABOLIC PROCESS
ATP METABOLIC PROCESS
MITOCHONDRIAL ELECTRON TRANSPORT NADH TO UBIQUINONE
RESPONSE TO HYDROGEN PEROXIDE
CELLULAR RESPIRATION
HYDROGEN PEROXIDE METABOLIC PROCESS
AEROBIC RESPIRATION
ELECTRON TRANSPORT CHAIN
RESPIRATORY ELECTRON TRANSPORT CHAIN
REACTIVE OXYGEN SPECIES METABOLIC PROCESS
OXIDATIVE PHOSPHORYLATION
ATP SYNTHESIS COUPLED ELECTRON TRANSPORT
-2.1 -2.0 -1.9
NES
Monocyte-Macrophage
NEUTROPHIL HOMEOSTASIS
CYTOKINE MEDIATED SIGNALING PATHWAY
RESPONSE TO BACTERIUM
NEURON APOPTOTIC PROCESS
RESPONSE TO TUMOR NECROSIS FACTOR
RESPONSE TO MOLECULE OF BACTERIAL ORIGIN
CELLULAR RESPONSE TO TOXIC SUBSTANCE
CELL CELL ADHESION VIA PLASMA MEMBRANE ADHESION MOLECULES
CELLULAR OXIDANT DETOXIFICATION
DETOXIFICATION
VISUAL BEHAVIOR
VASCULAR ENDOTHELIAL GROWTH FACTOR PRODUCTION
REACTIVE OXYGEN SPECIES METABOLIC PROCESS
POSITIVE REGULATION OF NEURON APOPTOTIC PROCESS
RELEASE OF CYTOCHROME C FROM MITOCHONDRIA
POSITIVE REGULATION OF SYNAPTIC TRANSMISSION
HYDROGEN PEROXIDE METABOLIC PROCESS
HYDROGEN PEROXIDE CATABOLIC PROCESS
POSITIVE REGULATION OF VASCULAR ENDOTHELIAL GROWTH FACTOR PRODUCTION
RESPONSE TO FIBROBLAST GROWTH FACTOR
-2 -1 0 1 2
NES
MiddleEB
CYTOPLASMIC TRANSLATION
PEPTIDE BIOSYNTHETIC PROCESS
1.8 1.9 2.0 2.1
NES
setSize 100 200 300 400 500
Adjusted P-value 0.00 0.01 0.02 0.03 0.04 0.05

**Figure S4. Mutation-intrinsic biological effects in MDS samples**
GSEA results for the pseudo-bulk paired comparison of cells classified as MUT and WT in MDS samples by SpliceUp.
